# Functional analysis of recurrent non-coding variants in human melanoma

**DOI:** 10.1101/2022.06.30.498319

**Authors:** Paula M. Godoy, Anna P. Zarov, Charles K. Kaufman

## Abstract

Small nucleotide variants in non-coding regions of the genome can alter transcriptional regulation, leading to changes in gene expression which can activate oncogenic gene regulatory networks. Melanoma is heavily burdened by non-coding variants, representing over 99% of total genetic variation, including the well-characterized TERT promoter mutation. However, the compendium of regulatory non-coding variants is likely still functionally under-characterized. We developed a pipeline to identify hotspots, i.e. recurrently mutated regions, in melanoma containing putatively functional non-coding somatic variants that are located within predicted melanoma-specific regulatory regions. We identified hundreds of statistically significant hotspots, including the hotspot containing the TERT promoter variants, and focused in on a hotspot in the promoter of CDC20. We found that variants in the promoter of CDC20, which putatively disrupt an ETS motif, lead to lower transcriptional activity in reporter assays. Using CRISPR/Cas9, we generated an indel in the CDC20 promoter in a human A375 melanoma cell line and observed decreased expression of *CDC20*, changes in migration capabilities, and an altered transcriptional state previously associated with neural crest transcriptional programs and melanoma initiation. Overall, our analysis prioritized several recurrent functional non-coding variants that, through downregulation of *CDC20*, led to perturbation of key melanoma phenotypes.

## INTRODUCTION

With the widespread availability of whole-genome sequencing and fewer discoveries of novel functional coding mutations, recent efforts have increasingly focused on identification and characterization of variants in the non-coding space of cancer genomes. Cis-regulatory variants (CRV) modulate transcription by altering the regulatory landscape of a gene, which in turn can lead to dysregulation of genes involved in cancer-driving pathways. Identifying CRVs of interest is therefore, generally, a three-step process: (1) identification of variants by whole-genome or targeted sequencing, (2) validation of variants through reporter assays and/or precise genome editing, (3) and characterization of the effect of the gene targeted by the CRV on tumorigenesis or cancer cell biology. For example, TERT promoter mutations were one of the earliest highly recurrent non-coding mutations identified in melanoma and are remarkable due to both a strong activating effect and prevalence in multiple cancers^1–3^. Present in ∼80% of cutaneous melanomas, the TERT promoter mutation creates a novel ETS motif that leads to binding of GABPA and derepression of TERT^4^. The full extent of TERT’s influence on tumorigenesis, particularly via this regulatory variant, is still emerging, including its canonical role on telomere maintenance^3, 5^. Beyond TERT promoter variants, few other CRVs have been identified and characterized in melanoma^1, 2, 6–10^. The next most common mutations in cutaneous melanoma are coding mutations in the MAPK pathway, predominantly BRAF^V600E/K^ and NRAS^Q61K^, as well as loss of key tumor suppressors like TP53, PTEN, and CDKN2A^1^^1, 12^, all with relatively clear canonical growth regulatory and proliferative functions.

Taking a more global view of gene expression, numerous RNA-sequencing studies, both from bulk and single-cell sources, have detected distinct transcriptional states in various melanoma populations^13–16^. For example, high levels of *MITF* are associated with a more proliferative/melanocytic state, while high levels of *AXL* are associated with an invasive state^13, 15, 17^. In between these two states is a stable intermediate state, that has been recently identified, and is orchestrated by a different set of transcription factors^13^. Additionally, a neural crest transcriptional program, present in the developmental precursors of melanocytes, is prominent in the first cells of melanoma^18, 19^. Aside from amplifications of MITF in 5-10% of melanomas, no other recurrent protein coding mutations have been associated with these distinct transcriptional subpopulations^11, 12^, leading to our hypothesis that CRVs could be a source of transcriptional dysregulation.

Guided by the threefold process described above, we leveraged whole genome sequencing of 183 melanomas from the International Cancer Genome Consortium and 69 melanoma-specific chromatin functional datasets to identify recurrent non-coding variants enriched in potentially functional enhancers/promoters. We validated several variants in the *CDC20* promoter which decrease *CDC20* promoter/enhancer-dependent reporter gene expression. We went further to genome engineer a promoter indel using CRISPR/Cas9 at this variant location in melanoma cell lines, which leads to decreased CDC20 expression, and then characterize the potential effects of the variant on cell viability, migration, and global gene expression changes, specifically in a subset of neural crest transcription factors.

## RESULTS

### Putative regulatory regions in melanoma are enriched for hotspot mutations

To identify recurrent non-coding mutations in human melanoma, we used variants called from whole genome sequencing (WGS) data from the International Cancer Genome Consortium (ICGC), the largest collection of WGS for melanoma to our knowledge, including 183 melanoma samples made up of 75 primary tumors, 93 metastases, and 15 human melanoma cell lines, as exome sequencing does not include full promoters or distal regulatory elements. The bulk of these tumors are cutaneous (140) but includes 35 acral and 8 mucosal melanomas. A total of 20,894,255 substitutions and 96,467 indels were identified from the ICGC Melanoma cohort^12^.

To refine our search space, we collated 69 previously published ChIP-seq and ATAC-seq datasets that were specifically performed on melanoma or melanocyte samples^18, 20–38^ (Supplemental Table 1). We reasoned these regions of the genome are more likely to bind transcription/chromatin factors and refer to them as putative melanoma regulatory regions (pMRRs). Genomic regions outside the pMRRs (red box, indicated by the lack of peak, Figure 1A) serve as an empirical null distribution but still have large numbers of recurrent mutations.

**Figure 1.**
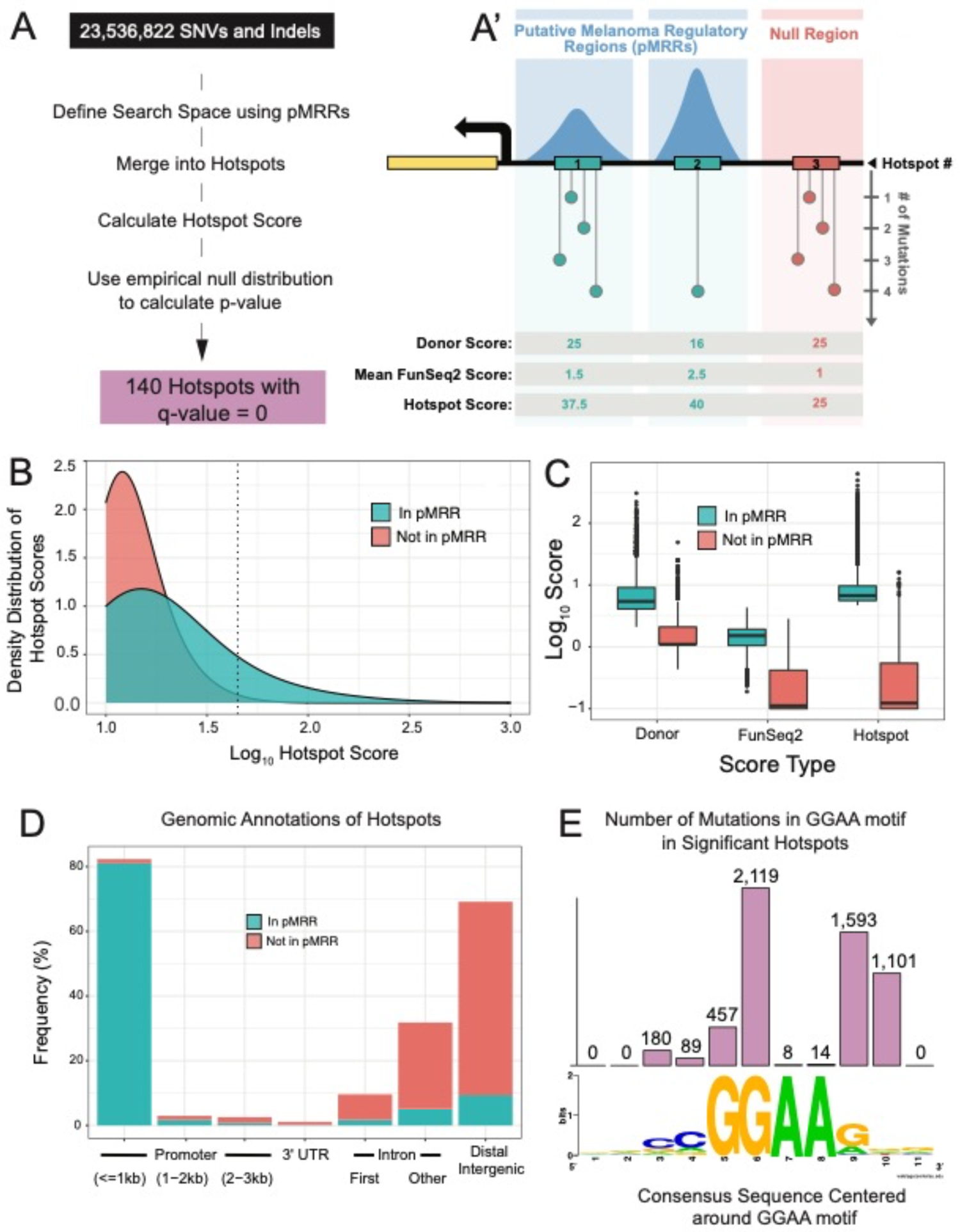
A method to identify putative functional non-coding variants in human melanoma. **(A)** Summary of pipeline to identify hotspots (A) with a generalized schematic of three theoretical hotspots (A’). Blue boxes indicate regions within putative Melanoma Regulatory Regions (pMRRs), and red box indicates null regions (i.e. those outside predicted regulatory regions). Numbered rectangles represent hotspots. Dot plots represent the number of variants within a given position. Donor score is equal to the square of the number of donors divided by the number of mutated positions, and FunSeq2 score is a weighting factor with higher values indicating higher conservation within regulatory regions and/or TF binding site motif altering. **(B)** Kernel density estimate of hotspot scores in pMRRs (blue) and not in pMRRs/in null regions (red). Hotspots with log_10_ scores lower than 1 are not shown. Dashed line depicts hotspot scores with a p-value = 1 x 10^-6^, lower p-values are to the right **(C)** Boxplots showing the log_10_-transformed Donor, FunSeq2, and Hotspot (Donor x FunSeq2) for the Top 10,000 highest-scoring hotspots. **(D)** Bar chart demonstrating the frequency of genomic annotations for Top 10,000 null hotspots (red bars) and statistically significant hotspots (707 hotspots, FDR-adjusted p-value < 0.05, blue bars). **(E)** Bar chart of the total number of mutations in significant hotspots (707 hotspots) at each site within 4 bp of the core ETS motif, GGAA (top, represents 5,561 mutations out of a total of 8,514 mutations), and WebLogo of 11 bp WT sequence (bottom).

pMRRs account for only ∼12% of the genome and harbor 2,142,063 variants (∼10% of total variants detected in the ICGC cohort). Of these, 444,161 variants are merged into 118,741 hotspots (3 or more variants within 25 bp are merged). Our empirical null distribution accounts for 5,478,131 variants within 1,462,992 hotspots. The remaining variants are isolated (i.e. not within 25 bp of another variant) and thus were not designated as hotspots.

All hotspots are also scored based on recurrence (donor score) and the average predicted impact of all variants within a hotspot as computed by the FunSeq2 algorithm, which weighs attributes such as evolutionary conservation and likelihood of TF motif creation/destruction (Funseq2 score, Figure 1A’)^39^. Hotspots in pMRRs have higher hotspot scores (product of donor score and FunSeq2 score) than those in null regions (Figure 1B). While donor scores are 4.9-fold higher in hotspots within pMRRs than those in null regions, FunSeq2 scores are 6.7-fold higher, drastically reducing the hotspot scores in regions outside of pMRRs and therefore potentially reducing false positives (Figure 1C).

Promoter regions are enriched in statistically significant test hotspots, while top-scoring null hotspots are commonly found in intergenic regions (Figure 1D). We identified 140 hotspots with FDR-adjusted p-values = 0 encompassing 2,631 mutations, notably including the known TERT promoter variant which has the 13^th^ highest hotspot score (Supplemental Table 2).

In order to evaluate for enrichment of putative TF binding site motifs, we used Homer analysis of pMRRs which identified motifs for TFs known to play prominent roles in melanoma, including SOX10^18, 40, 41^ (p-value = 1 x 10^-4^^72^) and ETS family factors^42^ (Supplemental Figure 1A), as well the multifunctional chromatin regulator CTCF (p-value = 1 x 10^-6092^). However, pMRRs that encompassed statistically significant hotspots are only enriched in ETS motifs, as previously observed^43^ (Supplemental Figure 1A). No ETS factor motifs are enriched in the mutant sequences, suggesting that most mutations break ETS transcription factor motifs (Supplemental Figure 1A). We found an almost identical distribution of mutations around the canonical GGAA ETS motif within the significant hotspots identified in our pipeline as previously reported^43^ (Figure 1E).

To focus our efforts on a candidate(s) among the top scoring hotspots (i.e. those with scores higher than TERT, encompassing thirteen candidates), we looked for consistent changes in gene expression for the gene nearest the recurrent variants between different stages of melanomagenesis (Supplemental Figure 1B). We used RNA-sequencing from 4 studies to calculate the fold change of the genes nearest to the hotspot between primary and metastatic tumors (The Cancer Genome Atlas, TCGA-SKCM^11^ and ICGC-MELA^12^), nevi and melanoma (Kunz^44^), and hPSC-derived melanoblasts with (KO melanoblasts) and without (WT melanoblasts) deletions in key tumor suppressors (Baggiolini^45^, see Methods for description of samples). *CDC20* (gene associated with the 8^th^ highest-scoring hotspot) is consistently upregulated in expression between melanoma and nevi (Kunz) and the KO and WT melanoblasts (Baggiolini, Supplemental Figure 1B). We observe a small increase in metastatic tumors compared to primary tumors in the ICGC cohort and no change between primary and metastatic tumors in the TCGA. The only other log_2_ fold-change greater than 1 is seen in the ICGC cohort for *TERT* expression (increase in metastatic melanoma, Supplemental Figure 1B). Low levels of *RPL18A* (3^rd^ highest-scoring hotspot), *HNRPNUL1* (6^th^), and *CDC20* (8^th^) tumors have higher survival rates than tumors with high expression of these genes (Supplemental Figure 1C). Taking both differential gene expression and association with survival rates for those with melanoma into consideration, we specifically focus on characterizing the *CDC20* promoter in melanoma.

### Variants in the CDC20 promoter have different effects on its transcriptional regulatory activity

The CDC20 promoter is mutated in 39 of 183 donors in the ICGC dataset, all of which are skin cutaneous melanomas (27.9% of cutaneous melanoma). The most common single-nucleotide variants (SNVs) are at adjacent positions chr1:43,824,528 (G>A, hereinafter termed G528A, mutated in 10 donors) and chr1:43,824,529 (G>A, G529A, 16 donors) as well as a SNV at position chr1:43,824,525 (G>A, G525A, 4 donors) and a multi-nucleotide variant (MNV) at positions chr1:43,824,528-43,824,529 (GG>AA, GG528AA, 4 donors) and are located within an ETS motif (Figure 2A). While at adjacent positions, G528A and G529A have different FunSeq2 scores (second number) and Genomic Evolution Rate Profiling (GERP) scores (third number) reflecting different degrees of purifying selection^46^. G525A is located within the core ETS motif, at the position that is most often mutated when taking all variants within statistically significant hotspots into consideration (Figure 1E) but is not the most recurrent variant in the CDC20 promoter hotspot, occurring only in 4/39 donors. Like G528A, G525A has both a high FunSeq2 score and a high GERP score.

**Figure 2.**
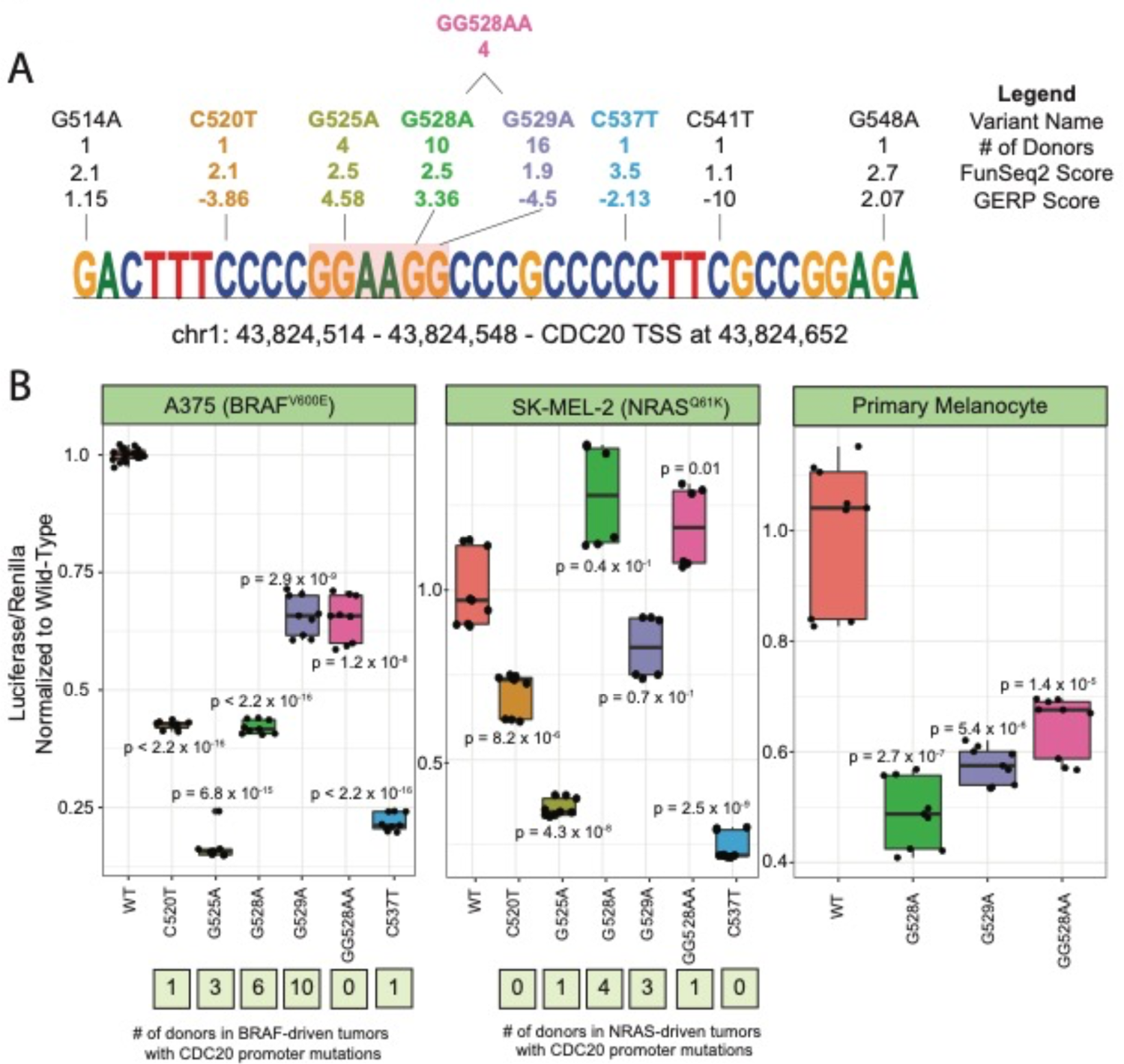
Functional analysis of recurrent CDC20 promoter variants. (A) The CDC20 promoter hotspot. All variants within the hotspot are denoted by name, # of donors with given mutation, FunSeq2 score, and GERP score. Co-occurring GG528AA double mutant is depicted above. Variants with colored text were validated by luciferase assay. **(B)** Altered CDC20 promoter activity for variants as assayed by luciferase reporter assays in melanoma (A375, SK-MEL5) and primary melanocytes. Boxplots depict normalized (to WT) luciferase assay results in these 3 different cell lines. The numbers below each boxplot indicate the number of donors in BRAF (left) or NRAS (right) tumors with the corresponding variant.

Overlaying chromatin-related assessments of the locus, the CDC20 promoter is accessible in 4/7 datasets that assay genome-wide chromatin accessibility (Supplemental Table 1). BRG1, CTCF, and TFAP2A are among the chromatin/transcription factors that have binding activity at the CDC20 promoter, as detected by ChIP-seq. ETV1, the only ETS factor with ChIP-seq data in our collation of melanoma-specific functional datasets (Supplemental Table 1), did not have binding activity at the CDC20 promoter in the 2 cell lines assayed (A375 and COLO-800, Supplemental Table 1).

To understand how the variants affect the regulatory activity of the CDC20 promoter, we performed luciferase assays using a 150 bp sequence length in a promoter-less luciferase vector. We assayed the 3 most prevalent variants, G528A, G529A, and GG528AA, in A375 (BRAF^V600E^), SK-MEL-2 (NRAS^Q61K^), and primary melanocytes (newborn foreskin melanocytes). We also assayed G525A, C520T, and C537T in A375 and SK-MEL-2. CDC20 promoter hotspots are not more likely to co-occur with pathogenic *BRAF* mutations than *NRAS* (p-value = 0.67, Fisher’s Exact Test, Supplemental Figure 3A).

Of the 4 variants present in more than one donor, G525A has the strongest effect on reporter activity, leading to a 6.1-fold decrease in A375 and 2.78-fold decrease in SK-MEL-2 (Figure 2B). G528A, G529A, and GG528AA, which flank the core ETS motif, lead to a decrease in reporter activity in A375 (2.4, 1.5, and 1.5-fold, respectively) and primary melanocytes (2.0, 1.7, 1.5-fold) but not in SK-MEL-2. Both G528A and G529A co-occur with BRAF and NRAS pathogenic mutations, but GG528AA is only detected with NRAS mutations in the available datasets. C520T and C537T, while only present in one donor each (BRAF only), also lead to a strong decrease in reporter activity (2.4 and 4.6 in A375, 1.4 and 4.1 in SK-MEL-2, respectively). We used motifBreakR^47^ to identify possible transcription factor binding motifs that are destroyed by the presence of the CDC20 promoter variants. As expected, the four variants closest to the core ETS motif are predicted to break sites for various ETS transcription factors, with G525A showing the largest reduction in ETS motifs of any variant (Supplemental Figure 2A).

We leveraged the RNA-sequencing data from a subset of the ICGC-MELA cohort to better predict the transcription factor (TF) that may have dysregulated binding in the mutated CDC20 promoter samples^12^. We reason that if TF_i_ binds to the CDC20 promoter at the core ETS motif that is disrupted by G525A, G528A, G529A, and GG528AA, *CDC20* expression will correlate with TF_i_ expression in WT samples but not in samples with the disrupted core ETS motif. Therefore, we calculated the Pearson correlation between every TF^48^ and *CDC20* in both WT and mutated samples. We identified 8 TFs that had high correlation of their expression (Pearson Correlation > 0.5) with *CDC20* expression in WT samples but low correlation in mutated samples (Supplemental Figure 2B). These include E2F1 and E2F2, which are known to regulate genes involved in cell cycle progression^49^ and, interestingly, the ETS family TF ELK1.

### CDC20-associated variants appear to be present as early clonal events but drop-out in distant metastatic melanomas

We next sought to understand the degree of clonality of the CDC20 promoter variants in sequenced tumors. Variant allele frequencies (VAF) can indicate how clonal a variant is by associating higher VAF with earlier appearance of the variant^50^. We compared the CDC20 promoter VAFs to those of the BRAF^V600E^, NRAS^Q61K/R^, and TERT G228A and G250A variants since these have all been reported to occur early^51^ (Figure 3A, Supplemental Figure 3B). In primary tumors, the BRAF, NRAS, and TERT variants are detected at median frequencies around 0.30, 0.32, and 0.41, respectively (Figure 3A). The median VAFs for the two most common CDC20 promoter variants G528A (0.34) and G529A (0.33) are only slightly lower than TERT and slightly higher than BRAF and NRAS, suggesting G528A and G529A mutations as early occurring events in melanomagenesis.

**Figure 3.**
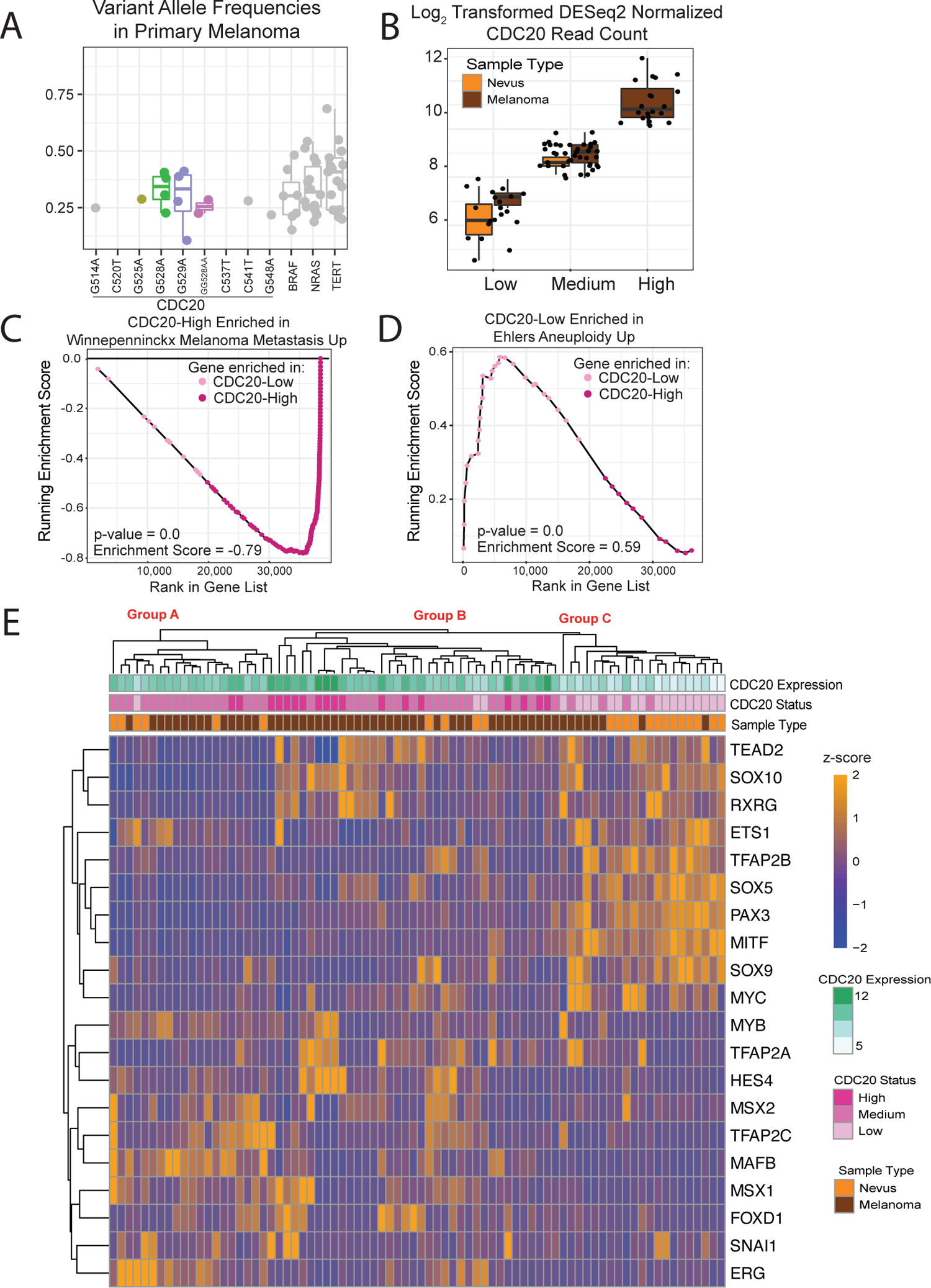
Changes in CDC20 expression levels correlate with specific gene expression programs. **(A)** Variant allele frequencies of the CDC20 promoter variants (each labelled), BRAF^V600E^ (BRAF), NRAS^Q61K^ and NRAS^Q61R^ (NRAS), TERT G228A and G250A (TERT) that are detected in primary melanomas. C520T and C541T are not detected in primary melanomas and have no data points. There are no statistically significant differences between G528A, G529A and other variants. **(B)** *CDC20* expression in CDC20-Low, CDC20-Medium, and CDC20-High nevus or melanoma samples from Kunz et al. Each data point represents the log_2_ DESeq2-normalized read count of *CDC20*. No nevi are classified as CDC20-high. **(C and D)** Gene set enrichment analysis of results for the Winnepenninckx Melanoma Metastasis Up gene set **(C)** and the Ehlers Aneuploidy Up gene set **(D)**. Each point represents a gene, ranked by expression, at the current running-sum statistic. Negative scores indicate enrichment in CDC20-high samples (as seen in C). Positive scores indicate enrichment in CDC20-low samples (as seen D). **(E)** Heatmap depicting z-score normalized expression patterns of 20 key neural crest transcription factors. Samples and genes are hierarchically clustered with orange and blue indicating relatively higher and lower gene expression, respectively, across samples. All columns are annotated by CDC20 expression (top row of boxes, log_2_ DESeq2-normalized read count), CDC20 expression group (second row, for low, medium, or high), and sample type (nevus in orange or melanoma in dark brown).

Since CDC20 promoter variants led to a decrease in reporter activity and CDC20 has been shown to be essential for migration in melanoma mouse models^52^, we hypothesized that promoter variants might decrease or disappear in later metastases. The G528A variant is detected mostly in lymph node metastases, often the first site of metastasis (n=6/11) and primary tumors (n=4/11). Only one distant metastatic sample out of a total of 51 had the G528A variant (Supplemental Figure 3C). Unlike G528A, G529A is detected across all stages and at median variant allele frequencies like those seen in earlier stages (Supplemental Figure 3B). Interestingly in A375 melanoma cells, G529A decreases reporter activity less than G528A, consistent with a model in which G529A, which is less deleterious to *CDC20* expression, does not seem to drop out in distant metastases like G528A, which lowers reporter expression more and, in agreement with published work^52^, thus would be disfavored in later metastases (Figure 2B).

### Distinct transcriptional programs emerge in nevi and melanoma in a CDC20 dosage-associated manner

To begin to pry into the differences between a CDC20-low and CDC20-high phenotype and how these differences may drive or support cancer progression at different stages of melanoma, especially those representative of the earliest states of melanoma, we utilized the Kunz cohort of 23 nevi and 57 primary melanomas that were RNA-sequenced^44^. We stratified all samples by *CDC20* expression with CDC20-high and CDC20-low classifications based on the 75^th^ and 25^th^ percentile of CDC20 expression, respectively. Remaining samples were classified as medium expression (Figure 3B).

We performed gene set enrichment analysis^53^ (GSEA) on CDC20-low and CDC20-high samples. As expected, based on prior studies, genes in the CDC20-high samples are enriched for gene sets associated with metastasis^52, 54, 55^(Figure 3C, Supplemental Table 3). CDC20-low samples have an enrichment of genes expressed in uveal melanomas with high aneuploidy^56^ (Figure 3D, Supplemental Table 2). This is in line with previous work that have shown increased aneuploidy in models with knockdown or mutated CDC20^57–59^.

We next sought to understand whether CDC20-low samples were enriched for key neural crest TFs, some of which (e.g. *SOX10*) are known to play important roles in melanoma initiation^18, 40, 41^. Hierarchical clustering using 20 neural crest TFs^60^ clustered samples into 3 major groups: samples with mostly low *CDC20* (Group C, median log_2_ expression = 6.7), samples with mostly high *CDC20* (Group B, median log_2_ expression = 9.4), and samples with medium *CDC20* expression (Group A, median log_2_ expression = 9.0, Figure 3E). This indicates differences in expression of these neural crest transcription factors is associated with *CDC20* expression. Surprisingly, CDC20-low samples cluster more closely with CDC20-high than CDC20-medium, despite having a larger difference in *CDC20* expression (Figure 3E).

Group C is made up of 13 nevi and 7 melanomas, 16 of which are classified as CDC20-low and four as CDC20-medium. This sample group has relatively high expression of genes prevalent in premigratory neural crest cells (*ETS1*, *SOX5*, *SOX9*, and *TFAP2B*) and melanocyte lineage specifiers (*SOX10* and *MITF*)^60^. Group A contains 6 nevi and 14 melanomas, 1 of which is classified as CDC20-low, 2 as CDC20-high, and 17 as CDC20-medium. This group has relatively high expression of *MYB* and *TFAP2B* which are prevalent in premigratory neural crest^60^, *MSX1* (neural plate border^60^), and *MAFB*, which is required for migrating cardiac neural crest cells^61^. Group B contains 32 melanomas and 4 nevi, 2 of which are classified as CDC20-low, 15 as CDC20-high, and 19 as medium. This group did not have relatively high expression across the group of any specific subset of transcription factors as seen with Group A and Group C. However, several isolated samples had relatively high expression of *SOX10*, *TEAD2*, and *RXRG*, as in Group C, and relatively high expression of *TFAP2A*, *MSX2*, and *HES4*, as in Group A. Notably, many of the neural crest transcription factors that are relatively higher in Group C are also known oncogenes in melanoma, particularly *SOX10*^18, 40, 62^, *MITF*^62^, *ETS1*^42^, and *MYC*^63^.

### Genome-engineered CDC20 promoter mutants have altered phenotypes and transcriptional profiles

Thus far, we have identified variants prevalent in the CDC20 promoter in melanoma tumors that by luciferase reporter assay reduce transcriptional activity and see distinct profiles of neural crest transcription factors in naturally occurring human melanoma tumors and nevi associated with high, medium, and low levels of *CDC20*^44^. To determine the effect of CDC20 promoter mutations on key cancer phenotypes and gene expression programs, we generated two CRISPR/Cas9-engineered A375 melanoma cell lines termed A3 and A10 (Figure 4A). The A3 line contains an indel on both alleles, both of which have the G528 and G529 nucleotides deleted. One allele retains the core GGAA motif while the other does not. The A10 line contains a larger deletion that completely removes the G525, G528, and G529 mutations, as well as the core ETS motif in both alleles (Figure 4A).

**Figure 4.**
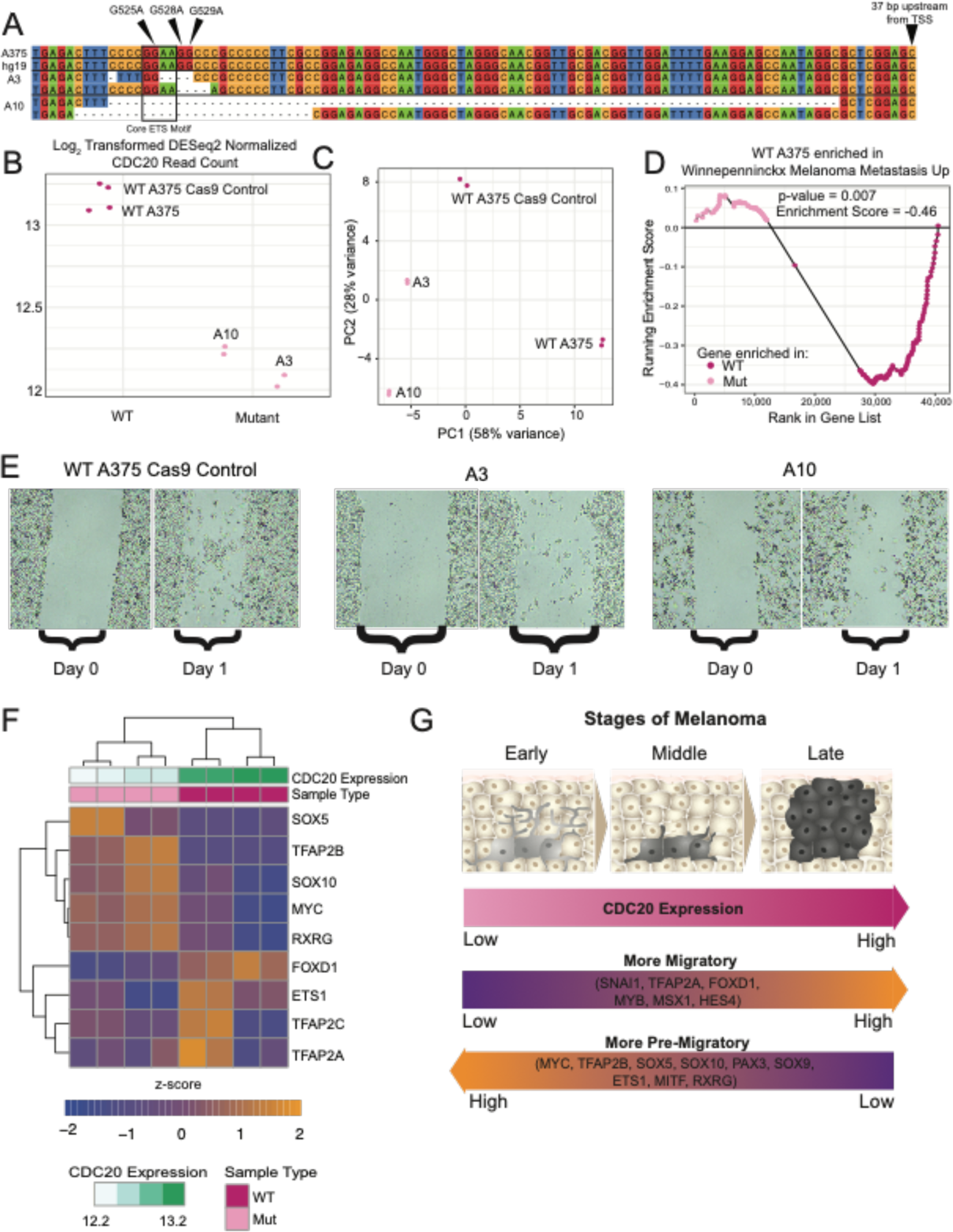
Engineered indels at the recurrently mutated CDC20 promoter locus leads to decreased CDC20 expression and changes in melanoma behavior. **(A)** Sequence alignment of the CDC20 promoter between hg19, WT A375, A3, and A10. Arrows denoting positions of G525A, G528A, and G529A. The ETS core motif is boxed. The last nucleotide of the sequence is 37 bp upstream of the TSS of *CDC20*. Nucleotides are color-coded and dashes indicate deletions. **(B)** Plot depicting log_2_ transformed DESeq2-normalized read counts of *CDC20* in WT A375 and CDC20 promoter indel strains, A3 and A10, with decreased CDC20 expression. Each point represents CDC20 expression in one sample. **(C)** Principal component analysis of read counts normalized by regularized log transformation using the top 500 most variable genes. The horizontal axis, PC1, explains 58% of the variance associated across all samples and separates out WT from CDC20 promoter indel cell lines. The vertical axis, PC2, explains 28% of the variance and separated A3 from A10. **(D)** Gene set enrichment analysis of results for the Winnepenninckx Melanoma Metastasis Up gene set. Each point represents a gene, ranked by expression, at the current running-sum statistic. Negative scores indicate enrichment in WT A375 samples as compared to the engineered indel lines A3 and A10. **(E)** CDC20 indel lines A3 and A10 show decreased migration capabilities compared to WT A375 cell lines. Images of scratch migration assay from day 0 (immediately after scratch) and day 1 (24 hours post-scratch). **(F)** Heatmap depicting z-score normalized expression patterns of 9 differentially expressed neural crest transcription factors. Samples and genes are hierarchically clustered with orange and blue indicating relatively higher or lower expression, respectively, of genes across samples. All columns are annotated by CDC20 expression (log_2_ DESeq2-normalized read count), and sample type (WT or mutant). *SOX5*, *TFAP2B*, *SOX10*, *MYC*, and *RXRG* are expressed at relatively higher levels in the CDC20 promoter indel-containing A3 and A10 cell lines. *FOXD1*, *ETS1*, *TFAP2C*, and *TFAP2A* are higher in WT (CDC20-high) A375 cell lines. **(G)** Model of *CDC20* expression and neural crest transcription factor signature over melanoma onset and progression. *CDC20* levels increase as melanoma progresses. Neural crest transcription factors that correlate with *CDC20* expression are more prevalent in migrating neural crest cells, whereas those that are relatively higher in CDC20-low settings are more prevalent in the melanocytic/pre-migratory neural crest states.

Both mutations decrease *CDC20* expression by 2.0-fold on average as detected by RNA-sequencing (FDR-adjusted p-value = 1.8 x 10^-40^, Figure 4B, Supplemental Table 4). The A3 strain has slightly lower *CDC20* expression than A10 despite having a smaller deletion and the retention of one core ETS motif (Figure 4B). Principal component analysis shows a separation along PC1 between the WT parental A375 line (high *CDC20*), the WT Cas9 control A375 line (high *CDC20*), and the mutant A3 and A10 line (low *CDC20*, Figure 4C).

Because CDC20 is an essential component of the cell cycle, we wondered if decreased *CDC20* levels would lead to decreases in cell viability^64^. We assayed viability in the presence of media containing serum, media containing serum and DMSO, and media containing serum and 30 nM dabrafenib (MAPKi) daily over the span of 6 days (Supplemental Figure 4A). A10 grows slightly slower than A375 and A3, despite having slightly higher levels of *CDC20* than A3 (Supplemental Figure 4A). No change in growth rates between A3 and A375 were observed (Supplemental Figure 4A).

We performed GSEA on the A375 WT and CDC20 promoter indel lines A3 and A10 using the same gene sets as above (Figure 3C and Figure 3D). There is significant enrichment of genes upregulated in WT vs CDC20 promoter indel cells for genes in the Winnepenninckx Melanoma Metastasis gene set^55^ (Figure 4D). While not statistically significant, we did see slight enrichment of genes upregulated in the mutant A375 line in the Ehlers aneuploidy gene set^56^ (Supplemental Table 3). To determine whether these gene sets are enriched in other melanoma cohorts, we performed GSEA on three other cohorts that underwent RNA-seq using the same *CDC20* stratification as before^11–13^ (Supplemental Table 3). All cohorts with high *CDC20* expression have statistically significant enrichment of genes associated with metastasis gene sets, while all CDC20-low cohorts except for TCGA-low have enrichment of genes in gene sets associated with low metastasis (Supplemental Table 3). The Kunz-low, TCGA-low, and Wouters-low samples have enrichment of genes in the Ehlers’ Aneuploidy gene set (Supplemental Table 3). Together these analyses show association of CDC20-high states with metastasis and CDC20-low states with aneuploidy across multiple cohorts.

To see whether our A375 promoter indel lines have altered migration capabilities as suggested by the results of GSEA and the literature^52^, we performed a scratch assay and observed decreased migration capabilities suggesting that, at least in this context, reduced levels of CDC20 affect migration more so than viability (Figure 4E). Because we see enrichment of an aneuploidy gene set in CDC20-low samples, we checked the A375 mutant lines for increased aneuploidy but did not observe any in a karyotyping analysis (Supplemental Figure 4B).

Finally, we performed hierarchical clustering using the 9 differentially expressed (FDR-adjusted p-value < 0.05) neural crest transcription factors from the 20 that were previously found to be correlated with differential CDC20 levels in naturally occurring nevi and melanomas (Figure 3E, Supplemental Table 4). We observe similar patterns of neural crest transcription factor levels in high and low CDC20 samples. *SOX10*, *SOX5*, *RXRG*, and *TFAP2B* are consistently high across the A375 mutant lines and CDC20-low samples in the Kunz cohort. *TFAP2A*, *TFAP2C*, and *FOXD1* were upregulated in WT A375 cells as well as in CDC20-medium or CDC20-high samples in the Kunz cohort.

Because we observe a similar association between neural crest transcription factor expression and *CDC20* expression between our CDC20 promoter indel cell lines and a cohort of nevi and melanoma, we looked for such patterns in the ICGC-MELA, TCGA-SKCM, and Wouters samples as well^11–13^. We stratified all samples by *CDC20* expression (low, medium, and high) and performed hierarchical clustering using the 20 neural crest transcription factors (Supplemental Figure 5A). We specifically looked at the 9 neural crest transcription factors that were differentially expressed between the CDC20 promoter indel and WT cell lines (Supplemental Figure 5B). *FOXD1* and *TFAP2C*, which are upregulated in WT lines compared to mutant, are also relatively highly expressed in CDC20-high or CDC20-medium samples in 3 out of the 4 cohorts. *TFAP2B*, *SOX5*, *RXRG*, and *MYC*, which are upregulated in the CDC20 promoter indel cells, are upregulated in CDC20-low or CDC20-medium samples in most or all cohorts. *TFAP2A*, *SOX10* and *ETS1* were relatively highly expressed in multiple CDC20-expression groups. In conclusion, we observe a consistent trend across 5 cohorts between certain neural crest transcription factors and *CDC20* expression.

## DISCUSSION

Using the largest available cohort of melanoma whole-genome sequencing data and several dozen melanoma-specific functional genomics datasets, we have identified hundreds of mutational hotspots containing putatively functional non-coding somatic variants. Under the assumption that variants outside of pMRRs are not, or are less likely to be, functional, we generated an empirical null distribution with which to calculate significance. We chose to focus on characterizing variants in the promoter of CDC20, whose weighted rate of recurrence and predicted functional significance were greater than that of the well-studied *TERT* promoter variants and began to investigate how these variants alter melanoma behavior.

At least half of the CDC20 promoter variants tested decreased reporter activity across all cell lines in this study. Four variants were within 2 bp of a core ETS motif but did not affect reporter activity to similar extents. G525A strongly reduced reporter activity in A375 and SK-MEL-2, while G528A, G529A, and GG528AA only reduced reporter activity in A375 and primary melanocytes, suggesting a cell-specific response to the variants. Most of these variants are not detected in distant metastases, which in agreement with previous work^52^, suggests that lower levels of *CDC20* may be disfavored in later metastases. Therefore, we propose a dosage-dependent role of *CDC20* on melanoma onset and progression, in which low levels of *CDC20* are important in early stages of melanoma but higher levels may be important for later stages of melanoma.

Like TERT, CDC20 performs a variety of canonical and non-canonical functions, many of which can be implicated in cancer formation^49, 52, 58, 59, 65–71^. Most crucial is its role in the cell cycle, where it interacts with the anaphase-promoting complex to degrade cyclin B and signal the end of metaphase and the start of anaphase^72^. Complete knock-down of *CDC20* is lethal but several studies have shown that partial knock-down or missense mutations that impair the ability of CDC20 to bind to other interacting proteins leads to aneuploidy, an important hallmark of cancer^57– 59, 73, 74^. Although aneuploidy did not increase in the CDC20 promoter indel A375 cell lines, CDC20-low samples in 3/5 cohorts analyzed had enrichment of genes associated with increased aneuploidy (Supplemental Table 3). Additionally, we only observed a slight reduction in growth rates in one CDC20 promoter indel strain, A10, which has similar levels of CDC20 as the A3 strain despite a larger deletion in the CDC20 promoter, suggesting that at least in our model, a 2-fold reduction of CDC20 does not significantly alter cell growth rates.

We have shown that three of the most common variants in the CDC20 promoter reduce reporter activity in a BRAF^V600E^-driven melanoma cell line and in primary melanocytes. Because we hypothesize that low *CDC20* levels are critical in early stages of melanoma, we looked for the enrichment of a neural crest transcriptional program, a state observed in the first malignant cells of melanoma^18, 40^. We show both in our genome engineered cells and in four other naturally occurring human melanoma cohorts that samples with low *CDC20* can have relatively high expression of certain neural crest transcription factors, such as *SOX10*, *RXRG*, *SOX5*, *TFAP2B*, and *MYC*. Notably, SOX10 and MYC have already been established as oncogenes^18, 40, 63^. RXRG reportedly drives a neural crest stem cell state that is resistant to treatment in a subset of cells in melanoma minimal residual disease^75^. TFAP2B regulates the melanocyte stem cell lineage in adult zebrafish^76^, and SOX5 has been shown to inhibit MITF^77^. Taken together, we hypothesize that low levels of CDC20 in melanocytic nevi may support a transcriptional program associated with a partially de-differentiated, neural-crest like state.

Meanwhile, as *CDC20* levels increase, we observe an increase in a different subset of neural crest transcription factors, including *FOXD1*, which is known to impair migration and invasion in melanoma models when knocked-down^78^. Therefore, as *CDC20* levels increase, cells may gain migration capabilities. In conjunction, we did not detect some of the more deleterious CDC20 promoter variants (i.e. those leading to lower reporter expression) in distant metastases. Additionally, we found enrichment of metastatic gene signatures in the CDC20-high expressing samples across all 5 melanoma cohorts (Supplemental Table 3) and observed loss of migratory capabilities in A3 and A10, the CDC20 promoter indel cell lines with lowered CDC20 levels.

The ever-expanding genomic data available for melanoma has been crucial in advancing our understanding of melanoma biology^11, 12^, but most of the largest datasets with publicly available clinical outcomes data (i.e. TCGA) overrepresent metastatic lesions, and even a subset of metastatic lesion types (i.e. lymph node metastases in TCGA). Thus, while *CDC20* has been implicated as a cancer-driving gene with higher levels often associated with melanoma metastases and poorer survival, we posit that specific levels of *CDC20* expression may be crucial to supporting or allowing passage of melanocytes through malignant transformation (*CDC20* low) to locally invasive cancer and then on to metastatic disease (*CDC20* high, Figure 4G). As in the case of MITF, a rheostat model of *CDC20* may exist, whereby higher levels of *CDC20* drives metastasis and lower levels support a phenotype likely beneficial in earlier tumors^79^.

## METHODS

### Calculating hotspot scores

*Step 1: Merge mutations into hotpots*. Mutation calls for SNVs and indels from the MELA-AU cohort were downloaded from dcc.icgc.org after receiving DACO approval^12^. Using a 25 bp window, we merged mutation calls using bedtools intersect^80^ into hotspots based on the premise that highly recurrent variants may be under positive selection at some point during the melanoma life cycle (e.g. favor melanoma growth) and that a transcription factor binding site(s) (TFBSs) may be disrupted/created by modifying any of multiple nucleotides in this window.

*Step 2: Filter hotspots not in putative enhancers/promoters.* We downloaded processed peak calls from ChIP-seq (e.g. H3K27Ac, H3K4me3, CTCF) and ATAC-Seq (revealing accessible chromatin domains) data from 69 melanoma datasets to enrich for putative Melanoma Regulatory Regions (pMRRs) which we reasoned are more likely to bind transcription/chromatin factors (Supplemental Table 1). These are indicated by the blue “peaks” in the example Figure 1A. We excluded exons and those regions (e.g. highly repetitive) from Encode excluded regions list^81^.

*Step 3: Calculate Donor Score.* The donor score for a given hotspot is represented as D^2^/G, where D is the number of samples (donors) with the specific variant and G is the number of nucleotide locations with variants in the hotspot. For example, in Figure 1A, the purple hotspot shows D = 3 + 1 + 2 + 4 = 10 mutations, at G = 4 different locations, for Donor Score of 10^2^/4 = 25.

*Step 4: Weight variants using FunSeq2 score*. Each mutation is weighted for predicted functional significance by features including predicted TFBS motif creating/breaking effect and evolutionary conservation using pre-computed scores from the published FunSeq2 algorithm (http://funseq2.gersteinlab.org/downloads) with a higher score predicting higher likelihood of functional significance^39^.

*Step 5: Calculate Hotspot Score.* Each hotspot is assigned a Hotspot Score as the product of the Donor Score (Step 3) and mean FunSeq2 score (Step 4) for all variants in the hotspot, to weigh both the number of variants and their predicted functional consequence in one metric. For example, in Figure 1A, the purple box shows (Average FunSeq2 score)*(Donor Score) = 1.5*25 = 37.5

*Step 6: Calculate p-value for each hotspot in MRRs relative to the empirical null distribution (non-pMRR regions from Step 2).* For each hotspot score within pMRRs, we calculated a p-value by determining the proportion of null hotspots with hotspot scores greater than or equal to it. All p-values were adjusted for false discovery rate (FDR). Adjusted p-values equal to 0 are provided (Supplemental Table 2).

### Genomic Analysis of Hotspots

For all pMRRs, statistically significant hotspots (FDR adjusted p-value < 0.05, 707 hotspots), and top-scoring hotspots outside of pMRRs (top 707 null hotspots by Hotspot Score), we annotated regions using the ChIPSeeker function *annotatePeak*^82^ (Figure 1D). For HOMER motif analysis, we ran findMotifsGenome.pl on BED files of all pMRRs and statistically significant hotspots to identify known motifs (Supplemental Figure 1A). For each variant within statistically significant hotspots, we made FASTA files with 20 bp sequences corresponding to either the WT or mutant sequence (variant at position 10). These were processed through HOMER using the findMotfs.pl function (Supplemental Figure 1A). A BED file containing only the CDC20 promoter variants were processed through motifBreakR^47^ using the known and discovered motif information from transcription factor ChIP-seq datasets in Encode^83^.

To calculate the ETS motif distribution, we first made FASTA files containing 11 bp sequences corresponding to either the WT or mutant sequence (variant at position 6) from the 707 statistically significant hotspots with FDR-adjusted p-values < 0.05. If a sequence contained the GGAA motif, we counted how far each variant within a statistically significant hotspot occurred from the nearest GGAA (if more than one instance was detected). If the reverse complement, TTCC was identified, as the nearest ETS motif, we first rewrote the sequence as its reverse complement and then counted the distance. A consensus sequence was generated with Web Logo (https://weblogo.berkeley.edu/logo.cgi) using a re-oriented version of the 11 bp WT fasta file where the first G of the GGAA motif is always at position 5.

### Cohort Comparison of Top 13 Genes

We downloaded DESeq2-normalized read counts from GSE112509 for the Kunz cohort and quantile-normalized read counts from Firehose (Broad GDAC) for the TCGA-SKCM cohort. The Kunz cohort is made of 23 laser-microdissected melanocytic nevi and 57 primary melanomas^44^. The TCGA cohort consists of 81 primary and 367 metastatic melanomas^11^.

For ICGC-MELA, we downloaded BAM outputs from STAR^84^ from the European Genome-Phenome Archive (EGA) under Study ID EGAD00001003353. Gene counts were calculated using RSEM^85^ and normalized by DESeq2^86^. This cohort consists of 56 melanomas from 46 donors and consists of 25 metastatic melanomas, 17 primary melanomas, and 14 cell lines derived from tumors^12^.

For the Baggiolini cohort, we obtained raw counts from the supplementary material of the corresponding publication and normalized counts by DESeq2^86^. This cohort is made up of human pluripotent stem cell derived cells that are engineered to contain doxycycline-inducible BRAF^V600E^. KO lines contain deletions to RB1, TP53, and P16. These cells were then differentiated into neural crest cells, melanoblasts, and melanocytes. For our study, we only considered WT and KO melanoblast samples that had activated BRAF^V600E^ expression. In line with the corresponding publication, we consider KO melanoblasts to be melanoma-like (based on the ability to form tumors when subcutaneously injected into NSG mice) while WT melanoblasts were considered to be a non-tumorigenic precursor to melanocytes^45^.

For each of the top 13 genes, we calculated the log_2_ fold-change between metastatic and primary melanomas (TCGA-SKCM and ICGC-MELA), primary melanoma and nevi (Kunz), and KO and WT melanoblasts.

Survival rates and corresponding p-values for high and low expressing tumors were downloaded from cBioPortal^87^ (TCGA-SKCM) using the Onco Query Language (OQL): *GENE*: EXP < −0.5 and GENE: EXP > 0.5. Data was downloaded from cBioPortal.org and plotted with ggplot2.

### Cell Culture

We obtained A375 cells from ATCC (CRL-1619). All A375s were grown in DMEM media (Corning, 10-013-CV) with 10% Fetal Bovine Serum (Gibco, 261470) and 1X Penicillin/Streptavidin (Pen/Strep, Sigma-Aldrich, P4333). SK-MEL-2 cells were obtained directly from the NCI-60 collection following written request and approval and were grown in RPMI-1640 media with 2 mM L-Glutamine (Gibco, 11875) with 10% FBS and 1X Pen/Strep. Newborn foreskin melanocytes were ordered from the specimen research core at the SPORE in Skin Cancer at Yale University. Primary melanocytes were grown in OPTI-MEM (Gibco, 31985) containing 5% FBS, 1X Pen/Strep, 10 ng bFGF (ConnStem, F1004), 4 mL of 5 mM IBMX (Sigma, #I-5879), 1 ng/mL Heparin (Sigma, #3393), and 200 µL of 0.1 M dbcAMP (Sigma, #D-0627). Cells were grown in a dedicated incubator set to 37°C at 5% CO2.

### Luciferase Assays

To make the luciferase vectors, we synthesized a 170 bp sequence containing the WT CDC20 promoter sequence (chr1:43,824,464-43,824,633) (GenScript). From this template, we amplified a 150 bp sequence using primers pGL3-CDC20_F and pGL3-CDC20_R (Phusion High-Fidelity PCR Master Mix, NEB M0531, Supplemental Table 5) that added restriction sites for SacI and XhoI to the 150 bp sequence. Both the pGL3-Basic Luciferase vector (Promega, E1751) and the CDC20 promoter amplicon were digested using SacI-HF (NEB, R3156S) and XhoI (NEB, R0146S) at 37°C overnight, followed by heat inactivation at 65°C for 20 minutes. Digested vector and amplicon were ligated using T4 DNA Ligase (NEB, M0202S) and transformed into OneShot Top10 Chemically Competent Cells (ThermoFisher, C404010). Individual colonies were mini-prepped and confirmed by Sanger Sequencing (Azenta).

Using the Q5 Site-Directed Mutagenesis kit (NEB, E0554), we induced variants in the WT sequence using primers designed by NEBaseChanger (https://nebasechanger.neb.com/, Supplemental Table 5). Sequences that were successfully mutated, as well as the WT pGL3-Basic vector and pRL-TK (Promega, E2241), were midi-prepped (Qiagen, 12941).

For A375, SK-MEL-2, and primary melanocyte transfections, 300,000 cells per well were seeded onto 6-well plates. All transfections were performed using 9 uL of Lipofectamine 2000 (Invitrogen, 11668), 1.5 µg of luciferase vector, and 1.0 µg of control pRL-TK (renilla), following the manufacturer’s protocol. All transfections were performed at minimum in duplicate.

The following day, luciferase and renilla luminescence were measured using the Dual-Luciferase Reporter Assay System (Promega, E1910) per manufacturer specifications. Cells were lysed using 500 µL of 1X Passive Lysis Buffer and incubated for 15 minutes on an orbital shaker. 20 µL of lysate were added to clear-bottom 96-well plates. We ran three technical replicates per sample. Luminescence was measured on a GloMax 96 Microplate Luminometer (Promega) using a standard Dual Reporter Assay program. All luciferase values were normalized to renilla, as the internal transfection control. We then normalized all variant ratios to the corresponding average WT value. p-values were calculated using Student’s t-test.

### Correlation between TFs and CDC20

The ICGC RNA-sequencing cohort consists of 56 samples from 46 donors. 13 samples (from 10 donors) contained a variant in the CDC20 promoter. The remaining 43 samples were WT samples. We downloaded a list of all transcription factors from http://humantfs.ccbr.utoronto.ca. We calculated the Pearson correlation between *CDC20* and every TF in either the samples with WT or mutant CDC20 promoter variants. We selected those that had correlations between the TF and the WT CDC20 promoter samples greater than 0.5 and less than 0.2 for those between the TF and mutant CDC20 promoter samples.

### Calculation of Variant Allele Frequencies

Mutation calls for SNVs and indels from the MELA-AU cohort were downloaded from dcc.icgc.org after receiving DACO approval^12^. We calculated variant allele frequencies by dividing the number of reads containing the variant divided by the total counts for each specimen. We then stratified specimens by tumor subtype (primary melanoma, lymph node metastasis, distant metastasis). VAFs from recurrent tumors or from cell lines derived from tumors were not considered. When plotting VAFs, the CDC20 promoter variants and the BRAF^V600E^ variants were plotted separately. Variant allele frequencies for NRAS^Q61K^ and NRAS^Q61R^ and for TERT G228A and G250A were combined into one boxplot each.

### Kunz RNA-sequencing analysis

We downloaded DESeq2-normalized read counts from GSE112509 for the Kunz cohort^44^. We classified each sample as CDC20-low, medium, or high based on *CDC20* expression. Samples with less than the 25^th^ percentile of *CDC20* expression were classified as low, while samples with greater than the 75^th^ percentile of *CDC20* expression were classified as high. All other samples are classified as having medium CDC20 expression.

We performed gene set enrichment analysis (GSEA) on the 20 CDC20-low and 20 CDC20-high samples using all gene sets in MSigDB (https://www.gsea-msigdb.org) that contained the keyword “melanoma”. We used the following parameters: 1000 permutations, the phenotypes were always set as low versus high (ergo enrichment scores are positive for CDC20-low, negative for CDC20-high), and permutations were performed on the gene set.

We manually curated a list of 20 neural crest transcription factors from two previously published sources^60, 88^. DESeq2-normalized read counts for these genes were used to construct the heatmap. Counts across every gene were scaled by setting the parameter “scale” to “row” in the heatmap plotting function *pheatmap.* Genes and samples were clustered using Euclidean distance.

### Genome Engineering of A375

A375 cells were nucleofected on a Lonza 4D nucleofector according to manufacturer recommendations (P3 solution, nucleofection program EH-100). Each nucleofection was performed with 1 x 10^5^ cells, 0.75 µL Cas9 Protein at 10 µg/µL (IDT v3 Cas9 protein, glycerol-free, # 10007806), and 0.75 µL of each sgRNA at 100uM (IDT) suspended in IDT Duplex Buffer (IDT, # 11-05-01-03) (Supplemental Table 5). Sham-nucleofections for WT A375 Cas9 controls were nucleofected with an equal volume of blank PBS. After nucleofection, cells were seeded into 500 µL of DMEM complete in a 24-well plate at standard incubator conditions.

72 hours post-nucleofection, cells were harvested, and split into 6-well culture for expansion and into lysis buffer for DNA extraction (homemade by GESC, formulation identical to Lucigen Quick-Extract buffer). PCRs were performed with Platinum Superfi II 2x master mix (Thermofisher, #12368010) and primers against the sgRNAs target site (Supplemental Table 5). PCR products were sequenced by NGS using Illumina.

After confirmation of cutting activity at, the pools were single-cell sorted using a Sony SH800 cell sorter at 1 cell per well into 4 x 96-well plates with 100uL of DMEM, with 50% conditioned media, 5 µM Rock Inhibitor, and 100 µM sodium pyruvate. Plates were allowed to grow for ∼10 days, then clones were harvested and re-screened using PCR primers against the targeted locus (Supplemental Table 5). Homozygous knockout clones were identified based on the presence of deletion junction and absence of the target locus. WT A375 Cas9 controls were sequenced at all gRNA target sites to confirm wild-type genotype. Homozygous knockout clones and wild-type Cas9 control clones were expanded, checked by STR profiling, tested for mycoplasma contamination, and used for subsequent experiments.

### Cell Viability Assay of A375 CDC20 Promoter Knock-outs and Controls

For each strain (A3, A10, and the wild-type Cas9 control), we seeded 1500 cells per well in a clear-bottom 96-well plate (Corning, #3903) in DMEM media containing 10% fetal bovine serum and 1X Pennicilin/Streptavidin (DMEM complete), DMEM complete with 30 nM dabrafenib (Selleck Chemicals, S2807), or DMEM complete with 1% DMSO. To measure viability, we used CellTiterGlo (Promega, G7570) as per the manufacturer’s protocol. Plates were read on a GloMax 96 Microplate Luminometer (Promega) using the standard CellTiterGlo program.

### Cell Migration Assay of A375 CDC20 Promoter Knock-outs and Controls

Scratch assays were performed by seeding 1 million cells per well in a 6-well plate in DMEM complete media. Using a P200 pipette, we scratched the plate at indicated positions. Cells were washed with 1X PBS and imaged on a Nikon Eclipse Ts2. Cells were then plated with DMEM media with 1% FBS and 1X Pen/Strep. On the following day, cells were washed with 1X PBS and imaged.

### RNA-sequencing of A375 CDC20 Promoter Knock-outs and Controls

300,000 cells of the parental A375 (in duplicate), two WT CRISPR/Cas9 clones (one replicate each), A3 (in duplicate), and A10 (in duplicate) were seeded on a 6-well plate. On the following day, we isolated RNA using the Qiagen RNeasy Plus Mini Kit (Qiagen, 74134). Samples were submitted to the Genome Technology Access Center at the McDonnell Genome Institute at Washington University School of Medicine for library preparation and sequencing.

Total RNA integrity was determined using Agilent Bioanalyzer or 4200 Tapestation. Library preparation was performed with 5 to 10ug of total RNA with a Bioanalyzer RIN score greater than 8.0. Ribosomal RNA was removed by poly-A selection using Oligo-dT beads (mRNA Direct kit, Life Technologies). mRNA was then fragmented in reverse transcriptase buffer and heating to 94 degrees for 8 minutes. mRNA was reverse transcribed to yield cDNA using SuperScript III RT enzyme (Life Technologies, per manufacturer’s instructions) and random hexamers. A second strand reaction was performed to yield ds-cDNA. cDNA was blunt ended, had an A base added to the 3’ ends, and then had Illumina sequencing adapters ligated to the ends. Ligated fragments were then amplified for 12-15 cycles using primers incorporating unique dual index tags. Fragments were sequenced on an Illumina NovaSeq-6000 using paired end reads extending 150 bases. RNA-seq reads were then aligned and quantitated to the Ensembl release 101 primary assembly with an Illumina DRAGEN Bio-IT on-premise server running version 3.9.3-8 software.

Read counts were normalized using DESeq2, comparing WT to mutant strains^86^. Principal component analysis was performed using the *plotPCA* function in the DESeq2 package. The heatmap was generated with *pheatmap* using z-score normalized counts of the manually curated list of 20 neural crest transcription factors^88^ with FDR-adjusted p-values < 0.1 (between WT and mutant samples).

Gene set enrichment analysis was performed as previously described using the 25^th^ and 75^th^ quantile to establish CDC20-low and CDC20-high expression groups, respectively. To generate heatmaps of all 4 cohorts, we downloaded Kunz, TCGA, and ICGC RNA-sequencing datasets as previously described. For the Wouters cohort, we downloaded normalized counts from bulk RNA-sequencing of 33 melanoma cultures^13^ (GSE134432). We calculated the mean across all samples classified as CDC20-low, medium, or high and plotted z-score normalized counts using the *pheatmap* function. Z-scores were calculated by scaling across rows, or genes.

### Karyotyping of A375 CDC20 Promoter Knock-outs and Controls

Karyotyping and analysis was performed at the Cytogenetics and Molecular Pathology Laboratory at Washington University School of Medicine. The cytogenetic test/ karyotype analysis was performed to assess aneuploidy (gains and losses of whole chromosomes), structural changes (chromosomal translocations, inversions, segmental deletions and duplications). This assay involves growing of cells in appropriate culture medium, hypotonic treatment, fixing cells, staining cells with GTG banding and microscopic examination. Twenty cells are counted for enumerating the number of chromosomes in a metaphase spread. Three of these metaphase spreads are digitally processed to produce a detailed karyotype/karyogram to perform a detailed study (analysis) for variant counts and structural aberrations. Analyzing a metaphase is defined by band-by-band comparison between chromosome pairs.

## DATA ACCESS

All raw and processed sequencing data generated in this study have been submitted to the NCBI Gene Expression Omnibus (GEO; https://www.ncbi.nlm.nih.gov/geo/) under accession number GSE206639.

## ACKNOWLEDGEMENTS

We thank the Genome Engineering and Stem Cell Center (GESC) at Washington University in St Louis for their Cell Line Engineering Services, the Cytogenetics and Molecular Pathology Laboratory at Washington University School of Medicine, and the Genome Technology Access Center at the McDonnell Genome Institute at Washington University School of Medicine for help with genomic analysis. The Genome Technology Access Center is partially supported by NCI Cancer Center Support Grant #P30 CA91842 to the Siteman Cancer Center from the National Center for Research Resources (NCRR), a component of the National Institutes of Health (NIH), and NIH Roadmap for Medical Research. This publication is solely the responsibility of the authors and does not necessarily represent the official view of NCRR or NIH. We thank Megan Glaeser and Rebecca Cunningham for critical reading of the manuscript and Catie Newsom-Stewart for help with figure conceptualization and design. This research was supported by the Melanoma Research Alliance Young Investigator Award #566840. P.G. was supported by NSF DGE-1745038.

## SUPPLEMENTAL FIGURE LEGENDS

**Supplemental Figure 1.**
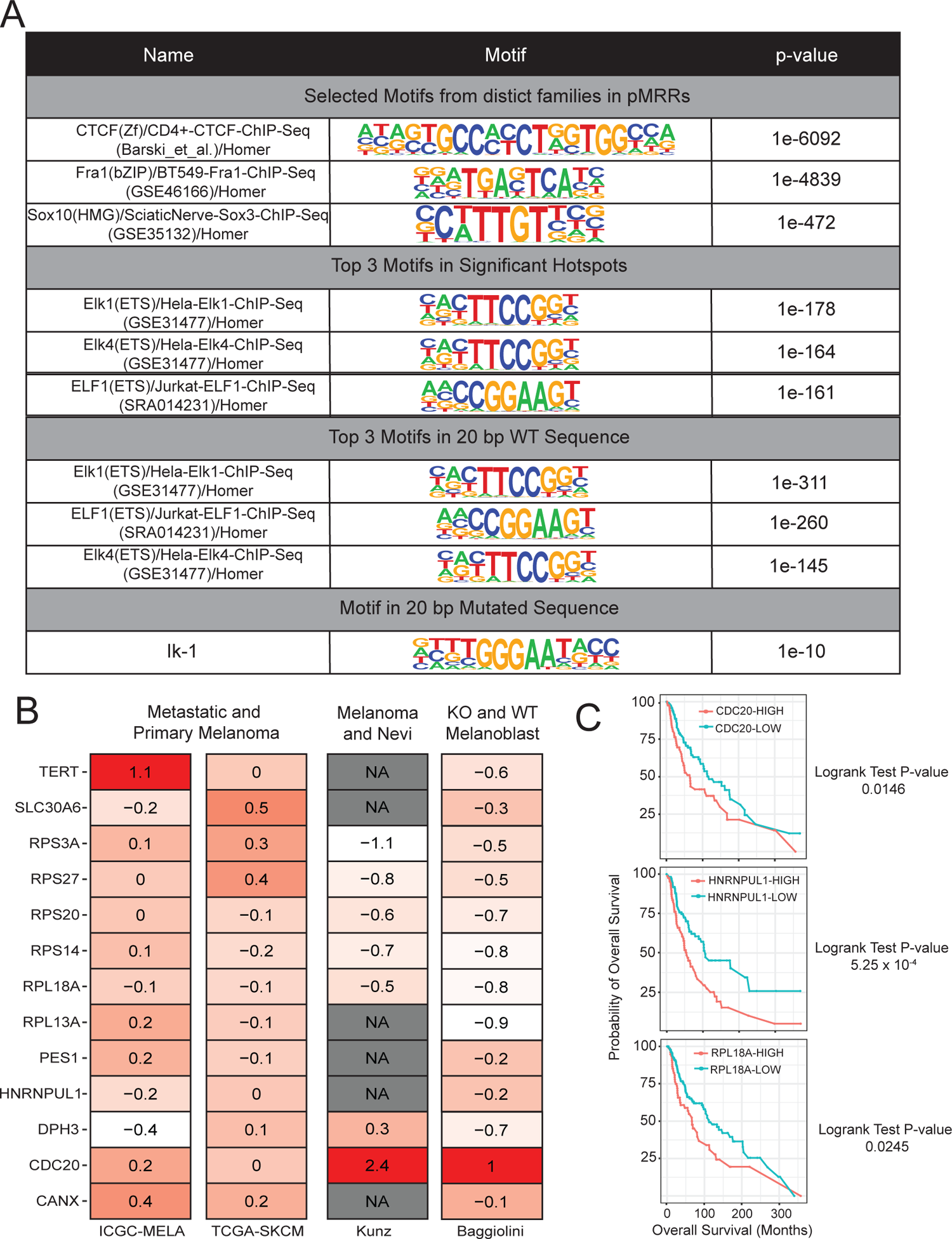
Characterization of putative melanoma regulatory regions, hotspots, and associated genes. **(A)** Table of selected motifs identified by Homer analysis. First section shows results for all pMRRs, regardless of whether region harbors a hotspot. To showcase diversity of transcription factors, we chose high-ranking motifs from three distinct transcriptional families. Second section shows top 3 motifs for pMRRs harboring statistically significant hotspots (707 hotspots, FDR-adjusted p-value < 0.05). Last two sections show top 3 motifs when input is a 20 bp sequence containing either the WT (top) or mutant (bottom) allele for all variants within statistically significant hotspots. **(B)** Log_2_ Fold-Change for Top 13 genes in ICGC-MELA, TCGA-SKCM, Kunz, and Baggiolini. Order in which samples are written represents numerator and denominator (e.g. if higher in metastatic, positive fold-change). **(C)** Kaplan-Meier curves representing over-all survival rates for high (red) and low (blue) expressing tumors for the three genes listed). Data and p-values obtained from cBioPortal using OQL.

**Supplemental Figure 2.**
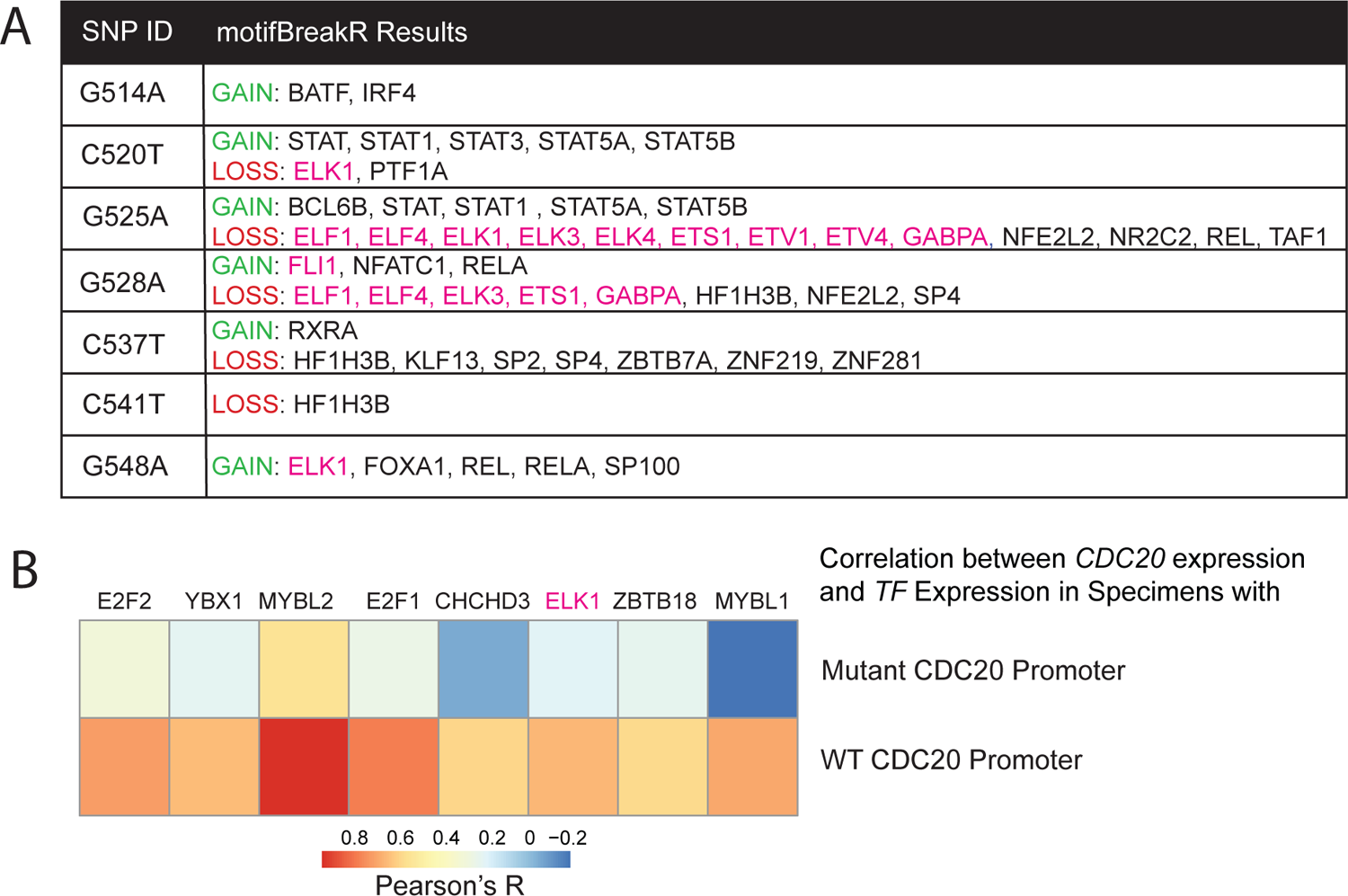
Motif impact of recurrent CDC20 promoter variants. **(A)** Table summarizing motifBreakR results. All transcription factors listed have strong and significant motif altering predictions. ETS transcription factors are colored in pink. **(B)** Heatmap of Pearson correlation values between TF (column) and samples with WT CDC20 promoters (top row) or mutant CDC20 promoters (bottom row.)

**Supplemental Figure 3.**
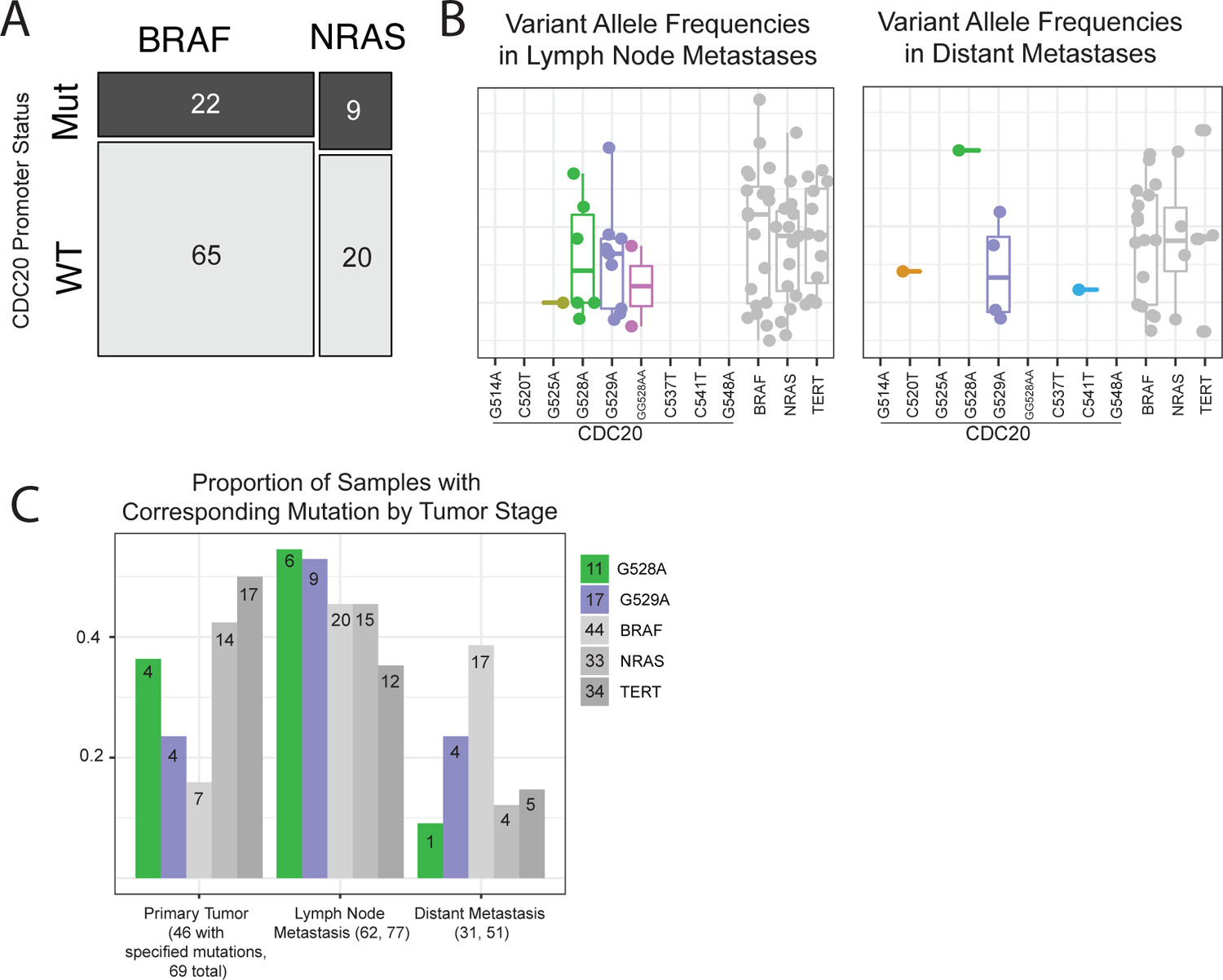
Co-occurrence and clonality of recurrent CDC20 promoter variants. **A)** Mosaic plot of the number of donors with either BRAF and/or NRAS mutations and WT and/or mutant CDC20 promoter. **(B)** Variant allele frequencies of the CDC20 promoter variants (each labelled), BRAF^V600E^ (BRAF), NRAS^Q61K^ and NRAS^Q61R^ (NRAS), TERT G228A and G250A (TERT) that are detected in lymph node metastases or distant metastases. **(C)** Each bar represents the number of specimens with the corresponding variant within the corresponding tumor subtype divided by the total number of specimens with the corresponding variant across all subtypes. Absolute counts for each subtype are labeled within the bar. Total counts across all subtypes are labeled within the legend. The total number of samples at each stage that contain one of the specified mutations are in parentheses below the row label; the number that follows is equal to the total number of samples in that tumor stage.

**Supplemental Figure 4.**
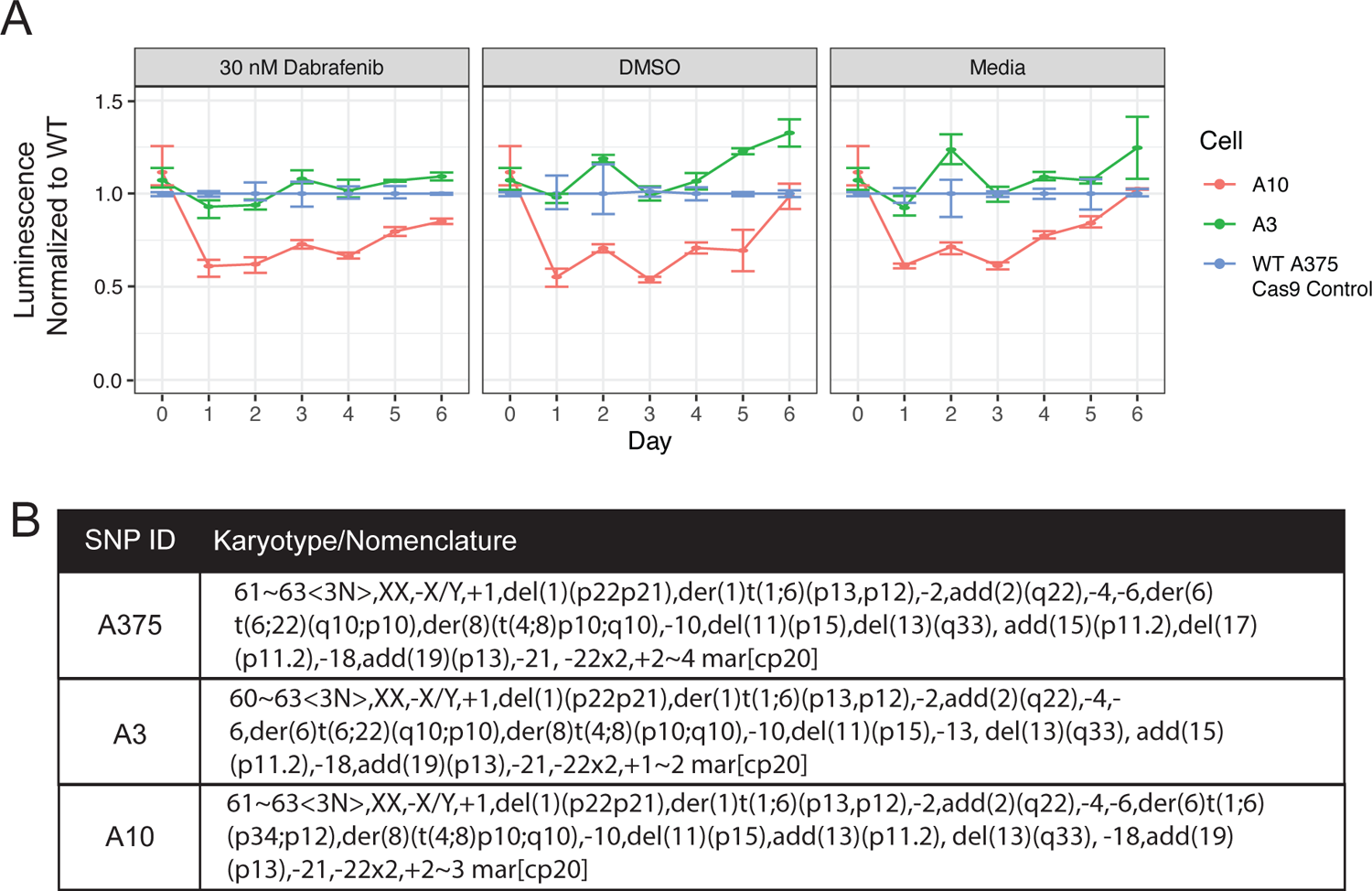
Viability and aneuploidy of WT and CDC20 promoter indel cell lines. (A) Proliferation rates are slightly lower in A10 but unchanged in A3. Plot shows luminescence values obtained from CellTiterGlo normalized to the average WT luminescence for each day and for each specific condition. Each point represents the average of three replicates. Confidence intervals are calculated using a nonparametric bootstrap method. **(B)** Table showing the karyotype/nomenclature of WT A375, A3, and A10. Differences across cell lines are color-coded.

**Supplemental Figure 5.**
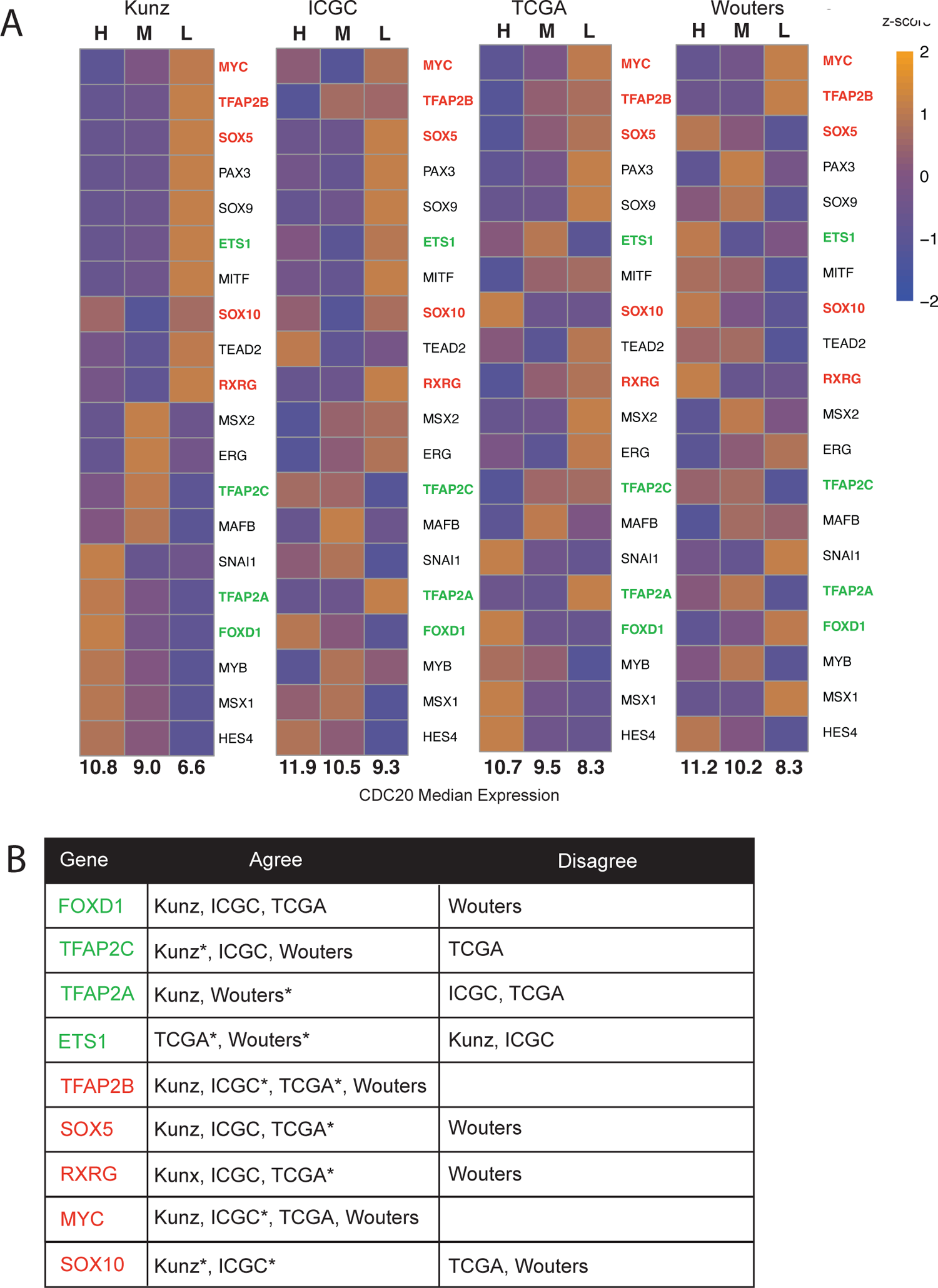
Neural crest transcription factor signature across 5 RNA-sequencing melanoma cohorts. (A) Heatmap depicting relative expression of 20 neural crest transcription factors using the average of all samples classified as CDC20-high, medium, or low. The median log_2_-normalized *CDC20* count is listed below each column of every heatmap. Orange indicates higher expression relative to other samples for the same gene. Genes in green are upregulated in WT A375 compared to CDC20 promoter indel cell lines. Genes in red are upregulated in CDC20 promoter indel cell lines. (B) Table summarizing whether a cohort has a relative gene level that matches or does not match the gene level seen in WT or CDC20 promoter indel cell lines. For a gene to agree, it needs to have relatively higher expression in the WT lines (green genes) or relatively higher expression in the CDC20 promoter indel lines (red genes). Cohorts that have an asterisk neither completely agree or disagree (e.g. relatively higher in CDC20-medium samples or relatively high in CDC20-low and CDC20-high, see SOX10 in TCGA).

